# Identification of germline chromatin modifying factors that influence zygotic transcription activation in *C. elegans*

**DOI:** 10.64898/2026.01.21.700918

**Authors:** Mariateresa Mazzetto, Paige Adekplor, Valerie Reinke

## Abstract

Precise control of the onset of transcription in post-fertilization embryos, often termed zygotic genome activation (ZGA), is essential to coordinate cell divisions and fate decisions to ensure proper development of the body plan. Maternally-contributed proteins and RNAs deposited in the zygote play an important role in ZGA, as does chromatin organization in pronuclei, yet the mechanisms by which parental germ cells anticipate the requirements for the first stages of ZGA is still not well understood. Using *C. elegans* as a model to study the epigenetic contributions from parental germ cells during the oocyte-to-embryo transition (OET), we previously demonstrated that histone modifications might act in germ cells to control ZGA. Here, we develop a novel assay to specifically track nascent transcription in gonads and embryos, which we use to identify chromatin regulatory factors acting in the germline that are responsible for subsequently promoting or inhibiting transcription during OET. This approach identifies regulation of both H3K4 methylation and H3K36 methylation in the parental germline as important contributors to successful ZGA and subsequent embryogenesis, and sets the stage for further mechanistic dissection of this critical developmental window upon the formation of a new organism.

## INTRODUCTION

The events surrounding gamete formation and function, fertilization, and early embryonic development are highly species-specific; indeed, these differences often underlie the inability of two similar species to cross-fertilize. Despite these fundamental differences, many common features are hallmarks of this critical developmental window. One shared feature is a period of transcriptional quiescence that initiates late in gametogenesis and persists through fertilization and the initial cell divisions of the newly formed embryo. Transcription then becomes reactivated at different rates and developmental stages in the embryos of various species, but this fundamental commonality suggests that a silent genome is critical for transition between generations. A major assumption is that silencing allows for reprogramming of the genome in the zygote to regain totipotency in preparation for development of a new organism. However, whether the underlying chromatin state inherited from the parents has any influence on genome reprogramming, or if only maternally provided cellular factors such as RNA and proteins drive reprogramming, is still poorly understood^1^. Here, we begin to address this question using the model organism *C. elegans*.

In the self-fertilizing hermaphrodite nematode *C. elegans*, transcriptional quiescence is initiated late in gametogenesis for both sperm and oocytes, through two mechanisms, one that directly prevents productive RNA Pol II elongation, and another that compacts chromosomes such that transcription is discouraged^2,3^. The genome remains silenced through fertilization, completion of meiosis, merging of two maternal and paternal pronuclei in the zygote, and the first two cell divisions, before active transcription is first detected in the four-cell stage. The genome becomes increasingly competent for gene expression, achieving a steady state around the 30-50 cell stage^4^. The *C. elegans* zygote is unique with regard to zygotic genome activation (ZGA), as the first division results in two cells with fundamentally distinct fates and lineages, AB and P, unlike most organisms, in which the first few zygotic cell divisions occur in the absence of fate specification. Thus, the genome first gains transcriptional competency in multiple cellular environments, and must already be prepared to turn on distinct gene expression programs from the very beginning. How the state of the parental chromatin is prepared to ensure both silencing and correct gene expression in the zygote under these circumstances is a fundamental question. We previously tracked histone modifications across the oocyte-to-embryo transition (OET) and found that H3K4me3 patterns are remodeled in developing oocytes primarily at a set of genes expressed at the initial onset of ZGA^5^. Notably, these genes primarily encode core regulatory proteins themselves, such as ribosomal proteins and RNA metabolic factors, suggesting that they are turned on first in preparation for processing of subsequent newly-transcribed factors later in embryogenesis. In addition, prior reports indicate that H3K27me3 and H3K36me3 patterns on parental gametes are important for establishing the subsequent germ lineage^6^, and a rapid turnover of the histone variant H3.3 also occurs concomitant with the first cell division^7^. These observations indicate that parental chromatin might indeed be important for the earliest activation of gene expression in the zygote. However, the mechanisms underlying these effects are not understood, and there are likely other chromatin-based mechanisms not yet identified that also contribute.

*C. elegans* provides an excellent model to address this issue, as the genome is compact and lacks extensive large-scale three-dimensional topology that complicates interpretation of chromatin state. Moreover, most regulatory information is proximal to gene promoters, making target gene identification straightforward. Finally, the oocyte-to-embryo transition (OET) is readily visible and accessible, aiding experimental assessment of chromatin and transcription. Therefore, we designed an assay using single molecule FISH^8^ that specifically tracks nascent RNA in gonads and early embryos using probes specific to introns. After confirming the specificity and accuracy of this reporter, we performed an initial analysis of a set of candidate regulators. This approach allowed us to further implicate H3K4me3 as well as H3K36me3, along with associated regulatory factors, in the proper silencing and reactivation of transcription across the OET, and sets the stage for further investigation into the chromatin-based mechanisms regulating this critical developmental stage.

## RESULTS

### Nascent RNA quantification in *C. elegans* gonads and embryos

To understand how changes in germline chromatin state in the parent might influence the initial onset of zygotic transcription, we established a method to distinguish newly transcribed, nascent mRNAs from the substantial population of maternally-provided processed transcripts loaded into the zygote. As a readout of nascent mRNA production, we employed single molecule fluorescent in situ hybridization (smFISH) because it reveals cellular and subcellular localization, as well as permits quantification of individual mRNA molecules^8^ (Figure 1A). Nascent RNA abundance can be uniquely tracked using probes designed to hybridize specifically to intron sequences, which are normally spliced out co-transcriptionally and rapidly degraded thereafter^9^. Multiple fluorescently-conjugated probes against a single transcript are necessary to reach the threshold of detection by confocal microscopy, so multiple large introns are needed to design sufficient overlapping probes for measurable signal, especially for short-lived introns. Notably, *C. elegans* genes often have relatively small and few introns compared to mammalian genes, reducing the number of genes that are good targets for nascent smFISH. Additionally, the reporter gene needs to be one of the very first mRNAs transcribed in the first wave of zygotic genome activation at the 4-cell stage. Our prior work had identified a set of genes characterized by two features: substantial remodeling of H3K4me3 in transcriptionally quiescent oocytes and early embryos, and sharply increased expression at the 4-cell stage in embryos^5^. Among this set, the *fust-1* gene had enough intron sequence that we could design probe sets using Stellaris program (Methods), sufficient to generate measurable fluorescent signal for smFISH (Figure 1B). *fust-1* exhibited substantial H3K4me3 remodeling during oogenesis, indicating that germline chromatin state might affect its early expression in embryos (Figure 1C). *fust-1* encodes a conserved, broadly expressed protein that binds RNA and has been implicated in splicing and miRNA regulation^10^.

**Figure 1.**
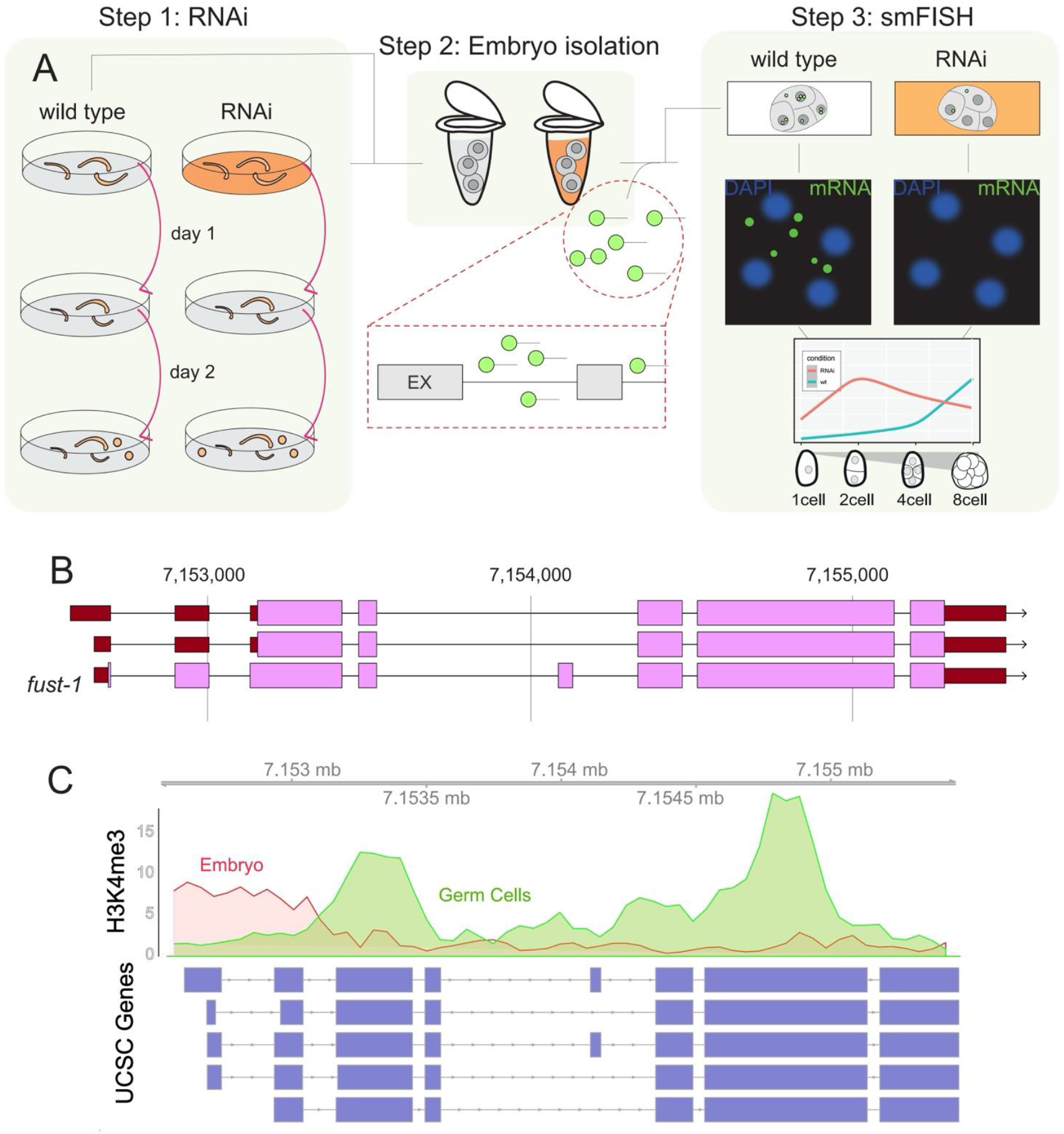
Experimental setup. A) Outline of smFISH protocol and its use as a reporter assay in RNAi screen of chromatin regulatory proteins. B) *fust-1* gene structure, highlighting introns and selected probes for smFISH. C) H3K4me3 remodeling at *fust-1* locus, demonstrating transition from broad domain across gene body to promoter-specific across OET.

To assess whether smFISH of *fust-1* introns provided a reliable and robust assay to measure nascent transcription in our hands, we tested several a priori expectations and performed control experiments. First, transcription is active in adult germ cells until oocyte differentiation commences, at which point several mechanisms converge and overlap to silence the genome through gamete maturation, fertilization, zygote formation, and the first two embryonic cell divisions^2,3^. At the 4-cell stage, there is an initial wave of transcription that becomes fully established by the 28-cell stage^4^. Thus, we expect that the intron-specific probes for *fust-1* should be detectable in the distal and medial germline and in 4-cell and later embryos, but not in oocytes or 1- or 2-cell stage embryos. smFISH of dissected gonads and early embryos confirmed this pattern (Figure 2A-B). We then tested two quantification methods (Aro^11^ and FishQuant^12^) to quantify signal abundance over this developmental timespan, confirming that we can accurately track and measure active *fust-1* transcription (Figure 2C).

**Figure 2.**
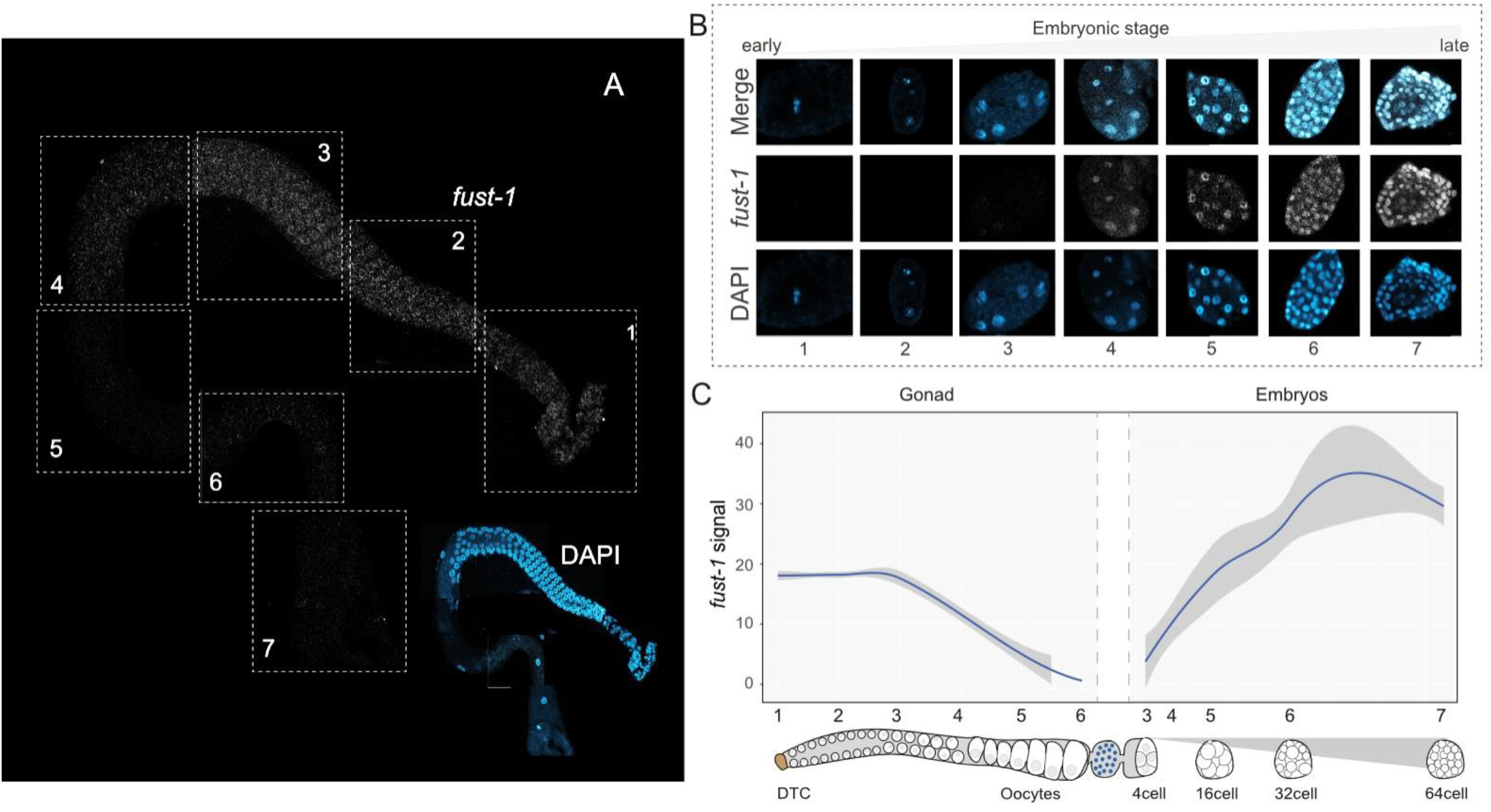
Establishment of intronic *fust-1* smFISH assay. A) Intronic *fust-1* smFISH in wild type dissected gonad, DAPI staining of same gonad in bottom right corner for reference. Boxes indicate sections of gonad for quantification in C). B) Intronic *fust-1* smFISH during embryonic development, from the 1 cell stage to the 60-100 cell stage. C). Quantification of gonad and embryonic signal in A and B, using BigFish/FISHQuant^12^.

A second expectation is that nascent, intron-retaining heteronuclear RNA should be preferentially detected in nuclei compared to the cytoplasm. Using DAPI staining to mark nuclei, we separately measured nuclear and cytoplasmic signal and found a three-fold increase in nuclear signal relative to the cytoplasm (Figure S1A). Additionally, we noticed concentration of *fust-1* signal in one to two subnuclear foci per nucleus that likely represent the *fust-1* genomic locus undergoing transcription (Figure S1B). We also separately designed exonic probes to *fust-1* (File S1), which should measure both nascent and mature *fust-1* transcripts. We then compared nuclear and cytoplasmic ratios for intronic and exonic probes, and demonstrated that the exonic probes were greatly enriched in the cytoplasm. In 2-cell embryos, only the exonic probes could be detected and were mostly cytoplasmic, demonstrating that a maternally provided transcript could be detected in transcriptionally quiescent embryos, even though the nascent transcript was not detectable (Figure S1C-E).

An additional expectation is that nascent (intronic) *fust-1* signal depends on active transcription. We therefore used the auxin-inducible degron (AID) system to acutely degrade an essential subunit of the RNA polymerase II complex, GTF-2H1^13^, in germ cells, followed by smFISH in early embryos. GTF-2H1-depleted embryos had essentially no *fust-1* intronic signal, while control (ethanol-treated) embryos displayed normal levels (Figure 3A, B). Finally, to confirm that the signal was specific to *fust-1*, we performed *fust-1(RNAi)* in adults, followed by *fust-1* intronic smFISH in embryos. Again, signal was significantly decreased when *fust-1* mRNAs were actively depleted by the RNAi pathway (Figure 3C). Altogether, these control experiments confirm that quantifying introns in *fust-1* transcripts by smFISH permits reliable, quantifiable assessment of nascent transcription in the adult gonad and early embryos. This reporter system should thus provide a robust, quantifiable assay to interrogate how changes to chromatin in the adult germ line result in altered initiation of zygotic genome activation in the early embryo.

**Figure 3.**
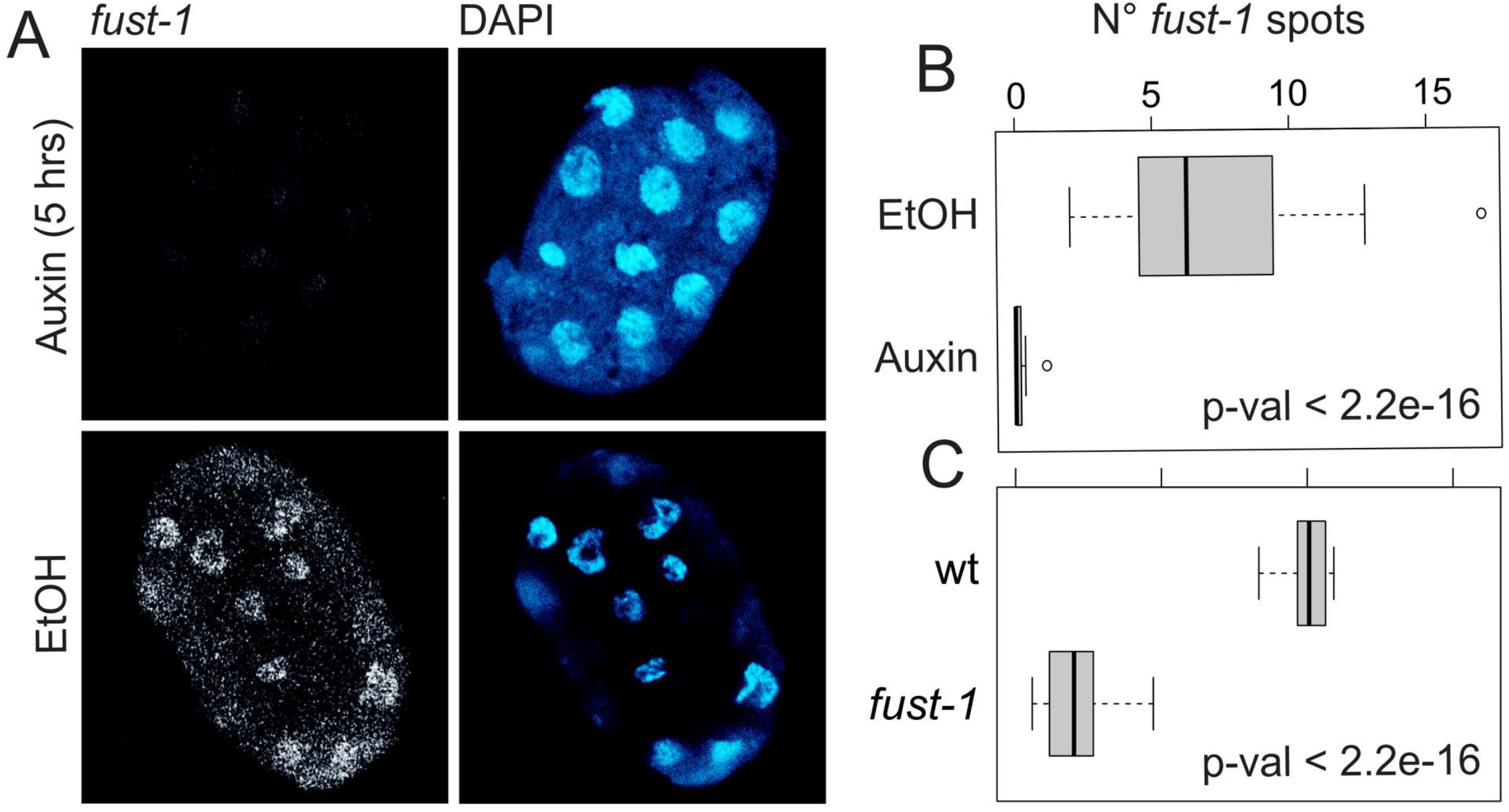
smFISH of nascent *fust-1* mRNAs depends on transcription of *fust-1*. A) No accumulation of *fust-1* nascent mRNA by smFISH upon induced degradation of GTF-2H1 by auxin treatment (5h) in parental gonad. B) Quantification of A. C) RNAi of *fust-1* results in decreased nascent *fust-1* mRNA in embryos.

### Screen for germline chromatin factors that might affect embryonic transcription

We therefore embarked on a preliminary screen of chromatin modifying proteins. A large set of ∼200 candidates was collected from two prior lists^14,15^ and prioritized by predicted roles, as well as by availability of the bacterial clone from our local RNAi library. For each candidate, we initiated RNAi by placing worms at the larval stage L4 onto plates containing bacteria expressing the dsRNA of the candidate gene. After 24-48 hours, gravid adults were collected, bleached, and embryos collected, fixed, and hybridized with intronic *fust-1* probes, then imaged and quantified across early embryonic development. To date, we performed RNAi of 16 candidate chromatin modifying factors (Table 1), of which several have provided significant changes in nascent *fust-1* transcription patterns in early embryos.

**Table 1.** Summary of functions and phenotypes of chromatin regulators screened.

| Group | Gene | Molecular Function | Mutant/RNAi Phenotype (Wormbase) |
| --- | --- | --- | --- |
| 1 | *rbr-2* | H3K4me demethylase | temp sens. germ cell mortality |
| 1 | *cdk-1* | cyclin-dep kinase | disrupted meiosis/mitosis; early Emb |
| 1 | *set-25* | H3K9 HMT | no phenotype, slow growth/ste with H3K9 HMT MET-2 |
| 1 | *set-17* | H3K4 HMT | pro-sperm gene cluster expression, suppresses spr-5 (HDM) Ste |
| 1 | *capg-1* | condensin subunit | mitosis/meiosis chromatid segregation, larval arrest w/ no germ cells |
| 1 | *fcp-1* | CTD phosphatase | limits pol II activity during oocyte maturation; Emb |
| 2 | *top-2* | topoisomerase | txn silencing in oocytes; maternal effect Emb |
| 2 | *hcp-6* | Centromere associated | ts allele causes Ste/Emb |
| 2 | *wdr-5.1* | H3K4me3 HMT (MLL), scaffold | ts Ste across generations, germ cell fate transformation, deposits H3K4me3 in embryos |
| 3 | *swsn-1* | SWI/SNF nucleosome remodeler | Emb/larval lethal |
| 3 | *cbp-1* | CBP/p300 HAT | promotes H3K27ac, Emb/larval lethal |
| 4 | *ash-2* | H3K4me3 HMT (MLL), scaffolding | promotes H3K4me2 in embryos, not germ cells |
| 4 | *set-2* | H3K4me3 HMT (MLL), SET1-like | ts Ste across generations, germ cell fate transformation |
| 4 | *rbbp-5* | H3K4me HMT (MLL), scaffolding | promotes H3K4me1/2/3, germ cell fate transformation |
| 4 | *set-26* | H3K4me3 reader | germ cell defects, partially redundant w/ set-9 |
| 5 | *jmjd-2* | H3K36/K9 me2/3 demethylase | slowed germ cell replication, increased meiotic DSBs |

To understand the relationship between these factors and their effects on *fust-1* transcription, we performed principal component analysis and identified five groups of factors, each with distinct *fust-1* zygotic transcription patterns (Figure 4A). Nascent *fust-1* levels during early embryogenesis, through the ∼60 cell stage, were graphed for each group to distinguish different trajectories for each group of candidate regulatory factors relative to control RNAi (orange and green profiles, respectively, in Figure 4B). Group 4, consisting of *ash-2, set-2, rbbp-5*, and *set-26*, exhibited a *fust-1* pattern very similar to control RNAi, whereas group 1, encompassing *rbr-2, cdk-1, set-25, set-17, capg-1* and *fcp-1*, consistently failed to accumulate *fust-1* nascent transcripts throughout early embryogenesis. *fust-1* nascent transcript accumulation was delayed and decreased overall in group 2 (*top-2, hcp-6*, and *wdr-5*). Finally, group 3 (*swsn-1, cbp-1*) exhibits a surprising pattern, with high levels in very early embryos, at a time when control embryos have not yet begun to transcribe *fust-1*; however, those levels dropped precipitously before climbing later in embryogenesis. Similarly, group 5 (*jmjd-2*) had high nascent *fust-1* levels in early embryos, although that signal was sustained across development. These last two groups thus appear to have a defect in transcriptional silencing that normally occurs during the oocyte-to-embryo transition.

**Figure 4.**
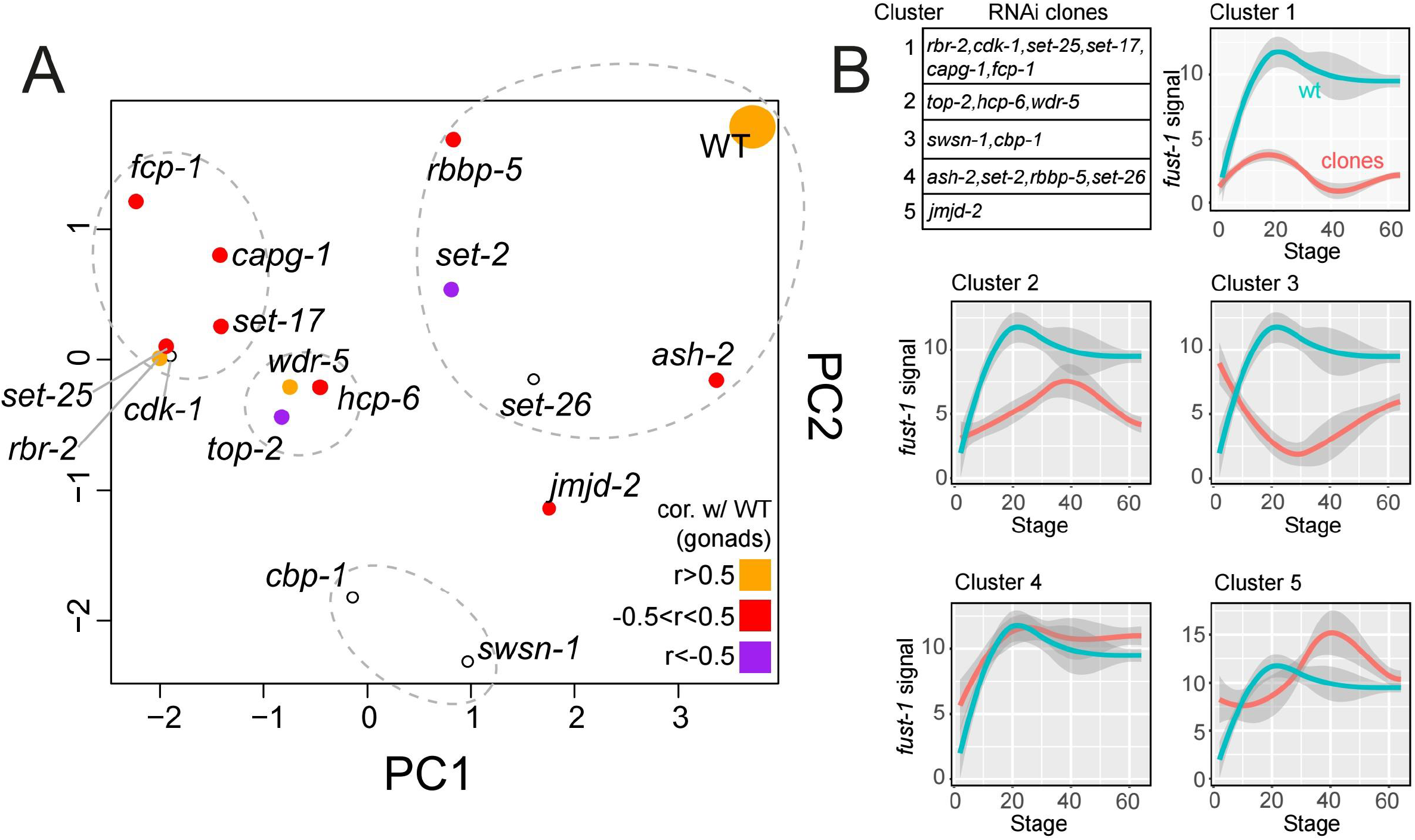
Identification of chromatin regulatory factors influencing *fust-1* zygotic transcription. Principal component analysis of the effects on embryonic *fust-1* transcription for the 16 factors fnctionally depleted by RNAi in parental gonads, identifying five functionally distinct groups. B) A table defining the candidate chromatin factors in each of the five clusters or groups, as well as graphs of the trajectory of *fust-1* transcription over early embryogenesis.

### Investigating relationships between candidate chromatin factors and nascent embryonic transcription

To follow up on these initial results, we developed several hypotheses for possible mechanisms underlying the changes in onset of *fust-1* transcription in the early embryo for candidate regulators in the different groups. We noted that several group 1 genes are required for early embryonic cell divisions, such as *cdk-1*^16^ and *capg-1*^17^, and thus RNAi of these factors might have indirectly failed to increase *fust-1* expression because profound disruption of embryonic development precludes zygotic genome activation. However, RNAi of other group 1 genes, such as *rbr-2, set-25*, or *set-17* do not significantly disrupt embryonic development, and might be directly involved in promoting *fust-1* expression in the early embryo. These three factors are directly involved in either adding or removing methyl groups from H3K4 and H3K9, suggesting these two modifications are important for *fust-1* activation, consistent with our prior observation that remodeling of H3K4 methylation is dynamic through the oocyte-to-embryo transition^5^.

Functional depletion of individual candidates in group 4 does not result in an obvious effect on embryonic *fust-1* transcription relative to controls. Surprisingly this group includes three of the core components of the MLL complex (*set-2, ash-2*, and *rbbp-5*), which is the primary H3K4 methylase complex in *C. elegans*, responsible for the majority of apparent H3K4 methylation at most development stages^18,19^. However, prior studies indicate that loss of any one component does not completely disrupt all functions of the other components, and that these subunits have unique roles independent of H3K4 methylation^20^. We therefore tested whether germline RNAi of two components simultaneously had an effect on *fust-1* transcription. Indeed, *set-2(RNAi)/ash-2(RNAi)* resulted in a complete loss of *fust-1* nascent RNA signal in early embryos (Figure 5 A,B), unlike RNAi of each individual gene. Moreover, the combined germline RNAi of both *set-2* and *ash-2* led to a slight increase in embryonic lethality from 2-5% to 15% (Figure 5C), suggesting that defects in H3K4 methylation might contribute to failure to properly initiate early transcription with subsequent downstream disruption of embryonic development.

**Figure 5.**
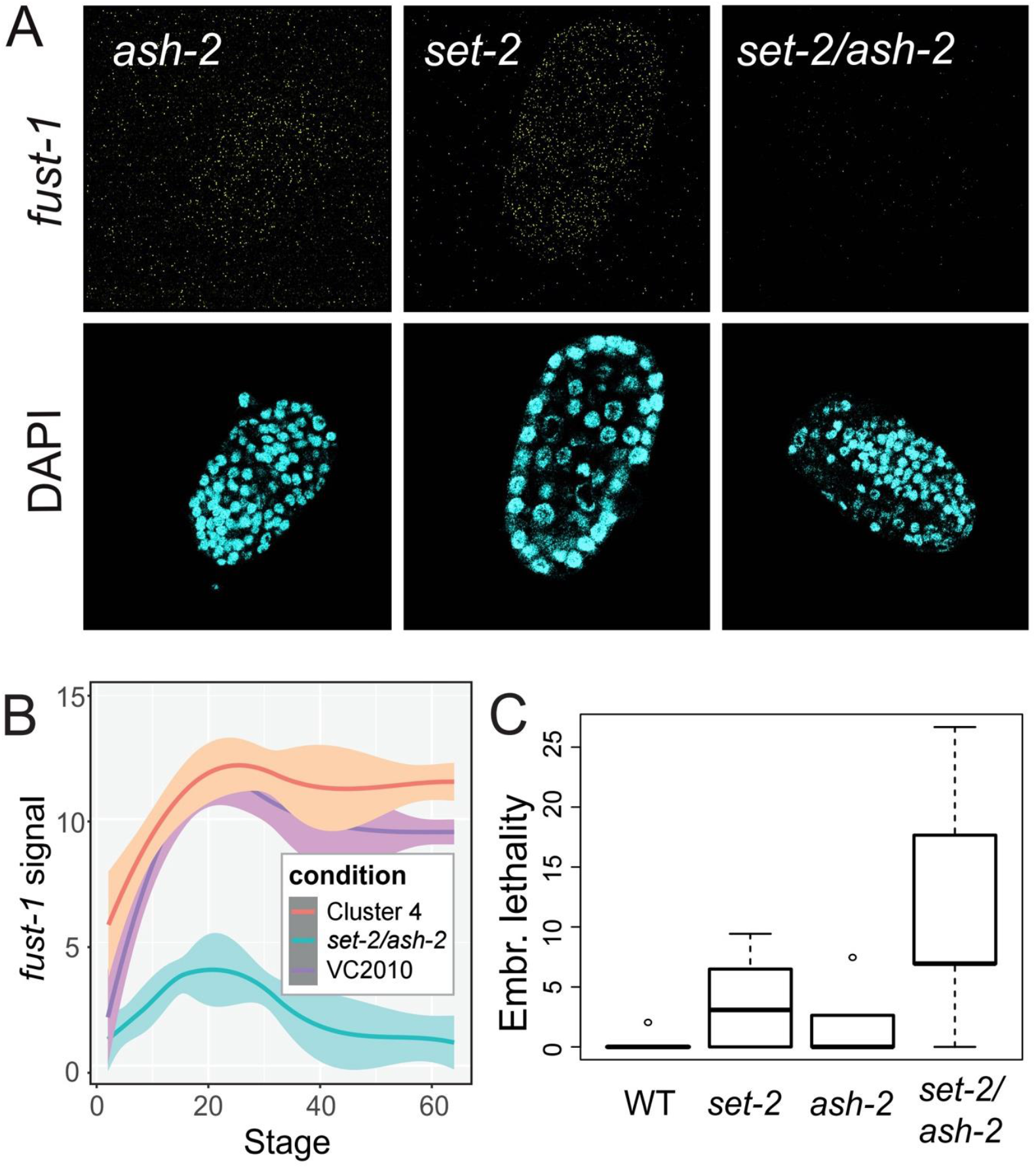
Combined depletion of MLL components disrupts *fust-1* transcription. A) RNAi of ash-2 and set-2 separately do not disrupt fust-1 smFISH signal, but combined RNAi of both factors results in complete loss of signal. B) Quantification of A. C) Extent of embryonic lethality for each RNAi condition.

Group 3 is characterized by an unexpectedly high level of nascent *fust-1* transcripts in very early embryos, which are normally transcriptionally quiescent, although these levels are not sustained as embryos continue to develop but fall well below the levels found in control RNAi treatments (Figure 4B). To determine whether the high levels of nascent *fust-1* in early embryos are due to a failure to silence transcription in oocytes, we performed smFISH in parental gonads treated with *cbp-1(RNAi)*, a group 3 candidate (Figure 6A). Indeed, nascent *fust-1* levels remain elevated in *cbp-1(RNAi)* oocytes, while in control oocytes, levels fall as transcription is inhibited. Notably, germline RNAi of *cbp-1* results in complete embryonic lethality (Figure 6B), which led us to investigate whether germline *cbp-1(RNAi)* also caused defects in germ cell differentiation and gamete formation. Adult gonads from *cbp-1(RNAi)*-treated animals displayed altered morphology, including expanded proliferation zones and delayed entry into oogenesis (Figure 6C), suggesting that this delay might result in aberrant or incomplete oocyte maturation, thus disrupting full implementation of transcriptional silencing during oogenesis. In *cbp-1(RNAi)* embryos, *fust-1* levels drop quickly as embryos develop, likely due to the severe defects in embryogenesis, although it is possible that failure to sustain transcription in in the early embryo also contributes to the severe phenotype. This observation is consistent with CBP-1 activity contributing to transcriptional silencing. However, *cbp-1* encodes a histone acetyltransferase responsible for promoting H3K27ac, a mark commonly found at promoters of actively expressed genes^21^. In wild type, levels of H3K27ac are high at promoters in isolated germ nuclei, but reduced substantially in early embryos (Figure 6D), suggesting that this mark, like H3K4me3, is remodeled during this developmental transition. Thus, whether changes to H3K27ac at promoters plays any direct role in transcriptional silencing during the oocyte-to-embryo transition is not clear.

**Figure 6.**
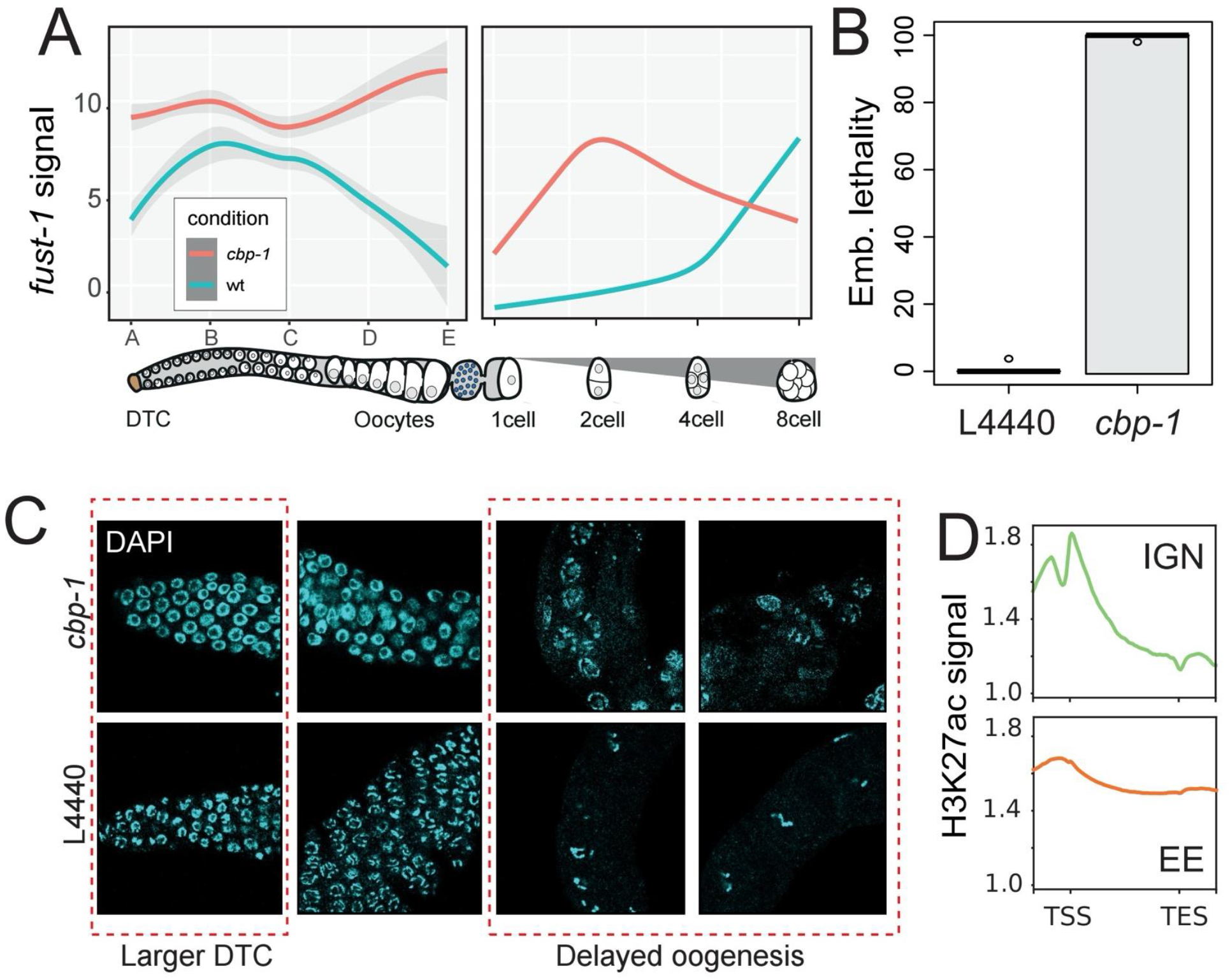
Histone acetyltransferase CBP-1 is required for normal oogenesis and appropriate transcriptional silencing of the genome. A) *cbp-1(RNAi)* results in elevated nascent *fust-1* transcription in both the parental gonad and the early embryo. B) *cbp-1(RNAi)* results in complete embryonic lethality. C) *cbp-1(RNAi)* results in defects of the onset of oogenesis. D) Relative H3K27ac levels in isolated germ nuclei (IGN) and embryos^5^ .

Finally, we investigated *jmjd-2*, the sole member of group 5, which is also characterized by high levels of nascent *fust-1* transcription in early embryos. This pattern differs from group 3 because *fust-1* transcript levels remain high as embryos develop, roughly to the extent of control embryos (Figure 4B). Notably, *jmjd-2(RNAi)* does not cause substantial embryonic lethality, which might result in indirect effects on *fust-1* transcription. Similar to *cbp-1*, however, *jmjd-2(RNAi)* results in elevated nascent *fust-1* levels in oocytes of treated animals (Figure 7A). *jmjd-2* is a member of the KMD4 family of lysine demethylases, and has been implicated in demethylation of both H3K36 and H3K9, an activating and repressive mark, respectively^22^. We therefore examined these two marks in wild type IGN and embryos (Figure 7B) and found that they exhibit opposite behaviors. While H3K9me3 levels become elevated in embryos, H3K36me3 levels drop, relative to germ cells. We hypothesized that JMJD-2 might be responsible for demethylating H3K36me3, leading to low levels in embryos. We examined H3K36me3 by immunostaining in embryos, and found higher levels in *jmjd-2(RNAi)* embryos (Figure 7C,D), particularly in very early embryos and in embryos at the second wave of zygotic genome activation (the 28-30 cell stage). Thus elevated H3K36me3 upon *jmjd-2(RNAi)* might promote ectopic transcription of *fust-1* and other genes. However, *jmjd-2(RNAi)* did not cause significant embryonic lethality, possibly because other regulatory events help to balance or rectify H3K36me3 levels as embryos develop, or elevated H3K36me3 is not sufficient to interrupt normal embryogenesis.

**Figure 7.**
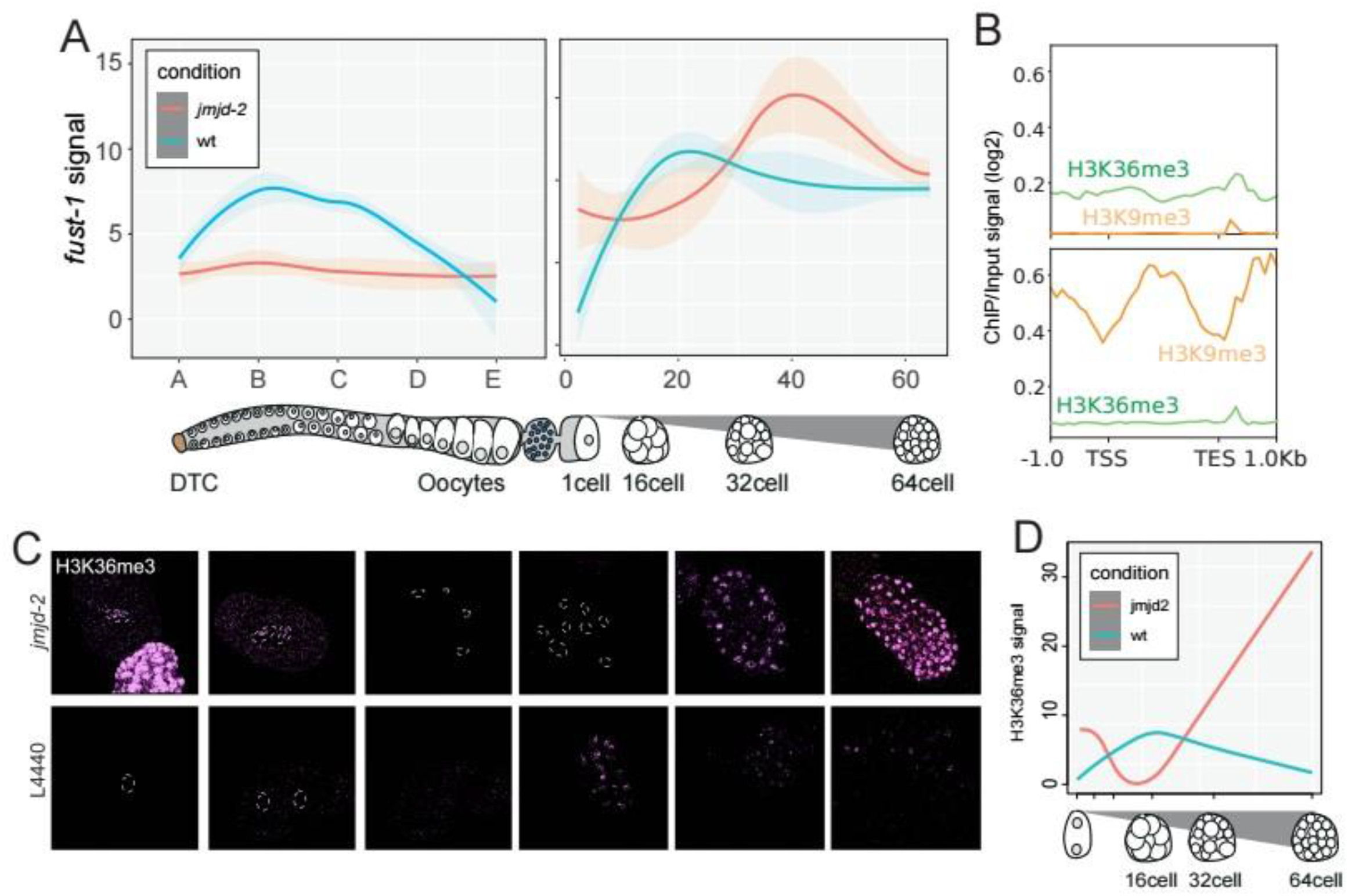
Lysine demethylase JMJD-2 is required for transcriptional silencing of the genome. A) *jmjd-2(RNAi)* results in elevated nascent *fust-1* transcription in both the parental gonad and the early embryo. B) Relative H3K36me3 and H3K9me3 profiles in IGN and embryos (ref). C) Immunostaining of H3K36me3 in embryos treated with either control (L4440) or *jmjd-2(RNAi)* showing elevated H3K36me3 in early embryos in the absence of *jmjd-2* function. D) Quantification of C.

## DISCUSSION

In this report, we describe development of a novel assay to follow the initial activation of transcription in early *C. elegans* embryos. Using smFISH, with probes specific to the introns of *fust-1*, allowed us to track nascent transcription in both gonads and early embryos. This strategy was capable of demarking the onset of genome silencing in developing oocytes, as well as the resumption of transcription in the early embryo, and permitted precise quantification of transcript abundance. We were then able to use this assay to read out whether functional depletion of chromatin regulatory factors in the parental germline had an effect on transcriptional dynamics across OET. The preliminary investigation of 16 factors identified several that disrupted these dynamics in distinct ways, including failing to completely inhibit transcription in oocytes, as well as failing to reinitiate transcription in 4-cell embryos. Thus, this report describes a new approach to investigating transcriptional regulation across OET.

Our initial follow up on selected factors from the 16 regulatory proteins investigated to date supported our previous findings that H3K4me3 might indeed play a role initiating zygotic genome activation. The highly conserved MLL complex is the primary factor that directs H3K4 methylation in *C. elegans*^20^. We found that dual disruption of two MLL components, *ash-2* and *set-2*, were sufficient to prevent zygotic expression of *fust-1*, although neither was able to do so individually. This result is consistent with previous observations that while these components are present in the MLL complex, they seem to have distinct and somewhat independent roles in promoting H3K4 methylation as well^20^. Other components such as *rbbp-5* and *wdr-5*.*1* that were included in our initial set, similarly had minimal or no obvious effects on *fust-1* zygotic transcription. Further investigation of how the MLL components together contribute to *fust-1* zygotic transcription, and the relationships between these components, is necessary.

Additionally, we followed up on *jmjd-2*, as RNAi of *jmjd-2* resulted in ongoing *fust-1* transcription in both oocytes and embryos, indicating a failure to induce transcriptional quiescence during OET. JMJD-2 encodes a KDM4-like demethylase that could potentially demethylate either or both H3K36 and H3K9, but when we examined existing histone modification profiles across germ cells and embryos, H3K36me3 levels dropped across this transition, while H3K9me3 levels rose. Given its role in demethylation, an effect on H3K36me3 seemed more likely. Indeed, staining against H3K36me3 in embryos indicated that indeed, loss of *jmjd-2* activity resulted in elevated H3K36me3 in 1- and 2-cell embryos in particular. However, the rate of embryonic lethality is relatively low upon *jmjd-2(RNAi)*, suggesting that this mis-regulation does not completely disrupt further embryonic development.

Overall, these results indicate that quantification of nascent RNA molecules via single molecule FISH is a versatile and information-rich reporter assay that leads to insights into potential mechanisms governing zygotic genome activation. Even testing a limited number of potential chromatin regulatory proteins, we were able to link the effects on nascent *fust-1* transcription to specific histone modification mechanisms that are temporally dynamic during the oocyte-to-embryo, setting the stage for deeper and wider investigation into chromatin-based contributions to the onset of zygotic gene expression.

## MATERIALS AND METHODS

### Worm growth and embryo preparation

*Caenorhabditis elegans* strains were maintained under standard conditions at 20 °C. For RNAi experiments, worms were synchronized by starvation and bleaching. Briefly, animals were chunked onto large NGM plates and allowed to starve for 4–5 days. Worms were floated to enrich for L1 larvae and plated onto large peptone-enriched plates to promote egg production. After 3 days, gravid adults were bleached, and embryos were allowed to hatch for 18–24 h before plating on large NGM plates. After ∼40 h, L4 larvae were transferred onto RNAi plates. To minimize escape from gene silencing, animals were transferred to fresh RNAi plates after 4–5 h. Embryos were collected 24 h after the second transfer.

For auxin-inducible degradation experiments, worms were grown following a dedicated auxin protocol. Briefly, synchronized worms were plated on peptone-enriched plates and bleached at the appropriate developmental stage. Auxin treatment was initiated ∼45 h after plating, following a 5-8 h pre-incubation (5 hrs for GTF-2H1 and 8 hrs for AMA-1) on both auxin and ethanol control plates. Embryos were collected once gravid adults contained 4–6 embryos.

### Gonad extraction

For gonad smFISH experiments, adults 16–20 h post-L4 were dissected to release gonads in 1.1x egg salts (10X egg salts: 250 mM HEPES pH 7.4, 1.18 M NaCl, 20 mM MgCl2, 20 mM CaCl2, 4.8 mM KCl) with 0.1% Tween 20 and 0.3 mM levamisole. Gonads were fixed with 3.7% formaldehyde, then immediately placed in −20°C methanol O/N and used the next day for hybridization.

### RNAi

RNA interference was performed by feeding using bacterial RNAi clones. RNAi plates were prepared in advance and allowed to dry for 5–7 days in the dark prior to seeding, followed by an additional 2 days of drying after bacterial seeding. L4 larvae were transferred onto RNAi plates and allowed to feed for 24–48 h before embryo collection. RNAi against these genes were used in this study: *wdr-5*.*1, rbbp-5, ash-2, set-2, set-26, set-17, swsn-1, rbr-2, cbp-1, cdk-1, top-2, hcp-6, smc-4, kle-2, capg-1, fcp-1, set-25, jmjd-2*. In addition, RNAi against *fust-1* served as a functional control.

### smFISH

#### a) smFISH probe design

smFISH probes were designed using the Stellaris RNA FISH Probe Designer (Biosearch Technologies). Probe sets targeted intronic regions of *fust-1* to specifically detect nascent transcripts. Probes were 17–22 bp in length, spaced by no more than 2 bp, and had a GC content of at least 25% (ideally ∼45%). Each probe set contained 32 probes and was labeled with a far-red fluorophore (Cy5/Quasar 670). Probes were stored dry at 2–8 °C in the dark and aliquoted upon resuspension.

#### b) Probe hybridization and staining

Embryos were collected by bleaching gravid adults and washed extensively with M9 buffer. Embryos were fixed in a fixation buffer (PBS, 3.7% formaldehyde, RNase-free water) for 15 min at room temperature, freeze-cracked in liquid nitrogen for 1 min, thawed, and incubated on ice for 20 min. Samples were washed twice in PBS and permeabilized overnight in 70% ethanol at 4 °C. For hybridization, embryos were washed in Wash Buffer A and incubated with hybridization buffer containing smFISH probes (final concentration 125 nM) for at least 4 h at 37 °C in the dark. Post-hybridization washes were performed using Wash Buffer A, followed by DAPI staining (5 ng/mL) at 37 °C. Samples were rinsed in Wash Buffer B and mounted in Vectashield.

#### c) Image acquisition and processing

Images were acquired using a Zeiss LSM 980 confocal microscope equipped with Airyscan 2. Typical acquisition settings included laser intensity of 0.18– 2%, exposure voltage of 850 V, and master gain of 0.2%. For datasets with elevated background signal, a duplicate image was generated and contrast-enhanced (100% contrast, −85% exposure) to suppress background and enhance true smFISH signal prior to quantification. This step was applied selectively and consistently across conditions.

#### d) Quantification and data analysis

smFISH signals were quantified using the Python implementation of FISH-QUANT. Individual fluorescent spots were identified, and both spot number and signal intensity were extracted. Nuclear and cytoplasmic signals were distinguished to confirm detection of nascent transcription. For each RNAi condition, transcriptional trajectories were compared to wild-type controls across embryonic stages. Log-transformed ratios of spot counts and intensity were calculated, and effects with |logFC| > 1 were considered significant. Principal component analysis (PCA) was used to classify RNAi conditions based on transcriptional dynamics.

### Embryo staining

For immunostaining experiments, *C. elegans* embryos were collected by washing gravid adults off multiple plates using M9 buffer. Worm suspensions were pooled into siliconized 15 mL or 50 mL conical tubes to prevent embryo loss and pelleted by centrifugation. Embryos were isolated by bleaching (7 mL for 15 mL tubes or 27.5 mL for 50 mL tubes) for 8–10 min, followed by vigorous vortexing and centrifugation at full speed for 40 s. Embryos were washed three times with M9 buffer, leaving ∼30 µL of buffer after the final wash.

Embryos were immediately flash-frozen in liquid nitrogen for 1 min and fixed by incubation in 1 mL of ice-cold 100% methanol for 20 min. Samples were centrifuged (900 rcf, 2 min), incubated in 1 mL of ice-cold 100% acetone for 10 min, and re-pelleted. Fixed embryos were blocked for 30 min at room temperature in PBS containing 1% BSA.

Primary antibodies were diluted in blocking buffer (typically 1:100–1:500) and incubated with embryos for 4 h at room temperature or overnight at 4 °C without agitation. Samples were washed once with PBS containing 5 ng/mL DAPI for 10 min with gentle rotation, followed by two washes in PBS (5 min each). Secondary antibodies were added in PBS (1–2 µL per mL) and incubated for at least 2 h at room temperature in the dark. Embryos were washed three times in PBS (10 min each), resuspended in 10% glycerol supplemented with antifade reagent, and mounted on slides (5–10 µL) for fluorescence microscopy.

### Brood Size and Embryonic Lethality analysis

To assess the impact of chromatin factor silencing on reproductive output and embryonic viability, adult worms were transferred to fresh RNAi plates 24 and 48 hours after the onset of RNAi treatment (referred to as Day 1 and Day 2, respectively). For each time point, brood size and embryonic lethality were quantified 24 hours after transfer. Brood size was determined by counting the total number of progeny, including both hatched larvae and unhatched embryos, present on each plate. Embryonic lethality was quantified by counting unhatched embryos, which were assumed to be non-viable. The percentage of embryonic lethality was calculated as the ratio of unhatched embryos to the total brood size.

## FIGURE LEGENDS

**Figure S1.**
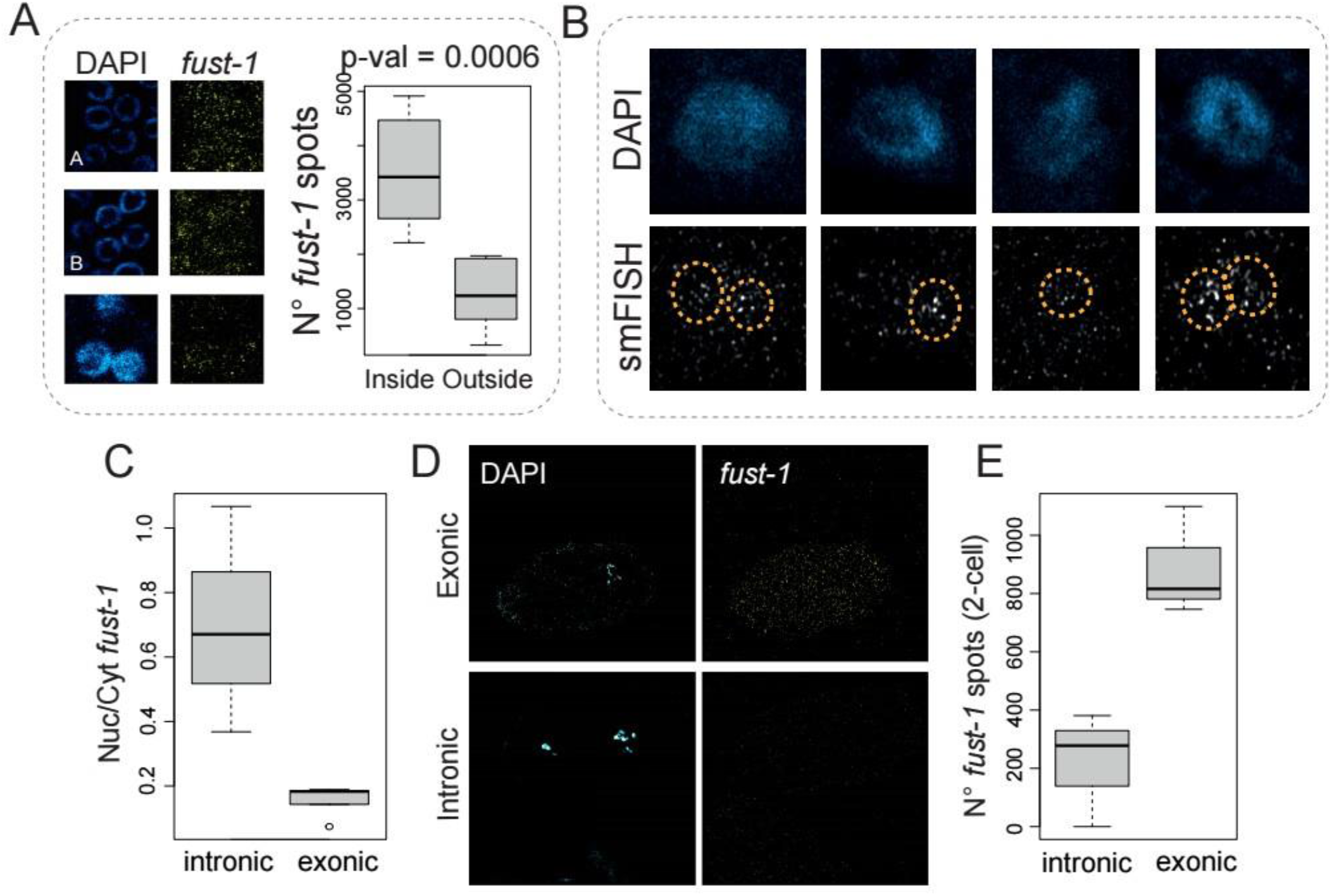
smFISH probes specific to *fust-1* introns are enriched in nuclei. A) smFISH in gonads, showing mitotic, early and late pachytene windows, including quantification of signal within and outside of nuclei. B) Intronic smFISH at the 4-cell stage of embryogenesis. C) *fust-1* smFISH using only exon probes was compared to only intron probes for relative abundance in the nucleus and cytoplasm, showing higher nuclear levels for intron probes. D) *fust-1* exon probes are detected in the 2-cell embryo where only mature, maternally-provided transcripts are present, whereas intron probes are not. E) Quantification of D.

